# A New World begomovirus infecting Cotton in Argentina

**DOI:** 10.1101/2020.02.20.957548

**Authors:** Verónica Delfosse, Humberto Debat, Diego Zavallo, Sabrina Moyano, Facundo Luna, Iván Bonacic Kresic, Sebastian Asurmendi, Sebastian Gomez-Talquenca, Ana Julia Distéfano

**Affiliations:** Instituto de Agrobiotecnología y Biología Molecular (IABIMO), Instituto Nacional de Tecnología Agropecuaria (INTA), Consejo Nacional de investigaciones Científicas y Tecnológicas (CONICET), Los Reseros y Nicolas Repeto, Hurlingham, Buenos Aires, Argentina, 1686; Instituto de Patología Vegetal, Centro de Investigaciones Agropecuarias, Instituto Nacional de Tecnología Agropecuaria (IPAVE-CIAP-INTA), 11 de setiembre 4755, Córdoba, Argentina, X5020ICA; Unidad de Fitopatología y Modelización Agrícola, Consejo Nacional de Investigaciones Científicas y Técnicas (UFYMA-CONICET), 11 de setiembre 4755, Córdoba, Argentina, X5020ICA; Estación Experimental Agropecuaria Mendoza, Instituto Nacional de Tecnología Agropecuaria (EEA-Mendoza-INTA), San Martín 3853, Luján de Cuyo, Mendoza, Argentina, 5534; Estación Experimental Agropecuaria Roque Saenz Peña, Instituto Nacional de Tecnología Agropecuaria (EEA-Saenz Peña-INTA), RN95 1108, Sáenz Peña, Chaco, Argentina

**Keywords:** Geminiviridae, Begomovirus, Cotton, New World Begomovirus, virus discovery, Gossypium

## Abstract

Cotton (*Gossypium* spp.) is a globally significant cash crop cultivated for its versatile fiber, widely used in the textile industry. Cotton, as other crops, is vulnerable to infectious pathogens. Several of them, including viruses, are a major threat to cotton production. Geminiviruses (family *Geminiviridae*) are insect transmitted, small non-enveloped viruses, with circular single-stranded DNA genomes, which are encapsidated in quasi-icosahedral geminated virions. Here we present evidence of a novel begomovirus (genus *Begomovirus*) infecting cotton from Argentina. Two circular ssDNA virus sequences were assembled from high-throughput sequencing data from *Gossyipium hirsutum* cotton samples showing mosaic symptoms from Argentina. Structural and functional annotation indicated that the virus sequences corresponded to complete DNA components A and B of a novel New World bipartite begomovirus. Genetic distance and evolutionary analyses support that the detected sequences correspond to a new virus, a tentative prototype member of a novel species which we propose the name “Cotton mosaic virus” (CoMV).

## Annotated Sequence Record

Cotton (*Gossypium* spp.) is an economically significant crop cultivated worldwide for its versatile fiber, which plays a pivotal role in the textile industry. However, the cotton industry faces numerous challenges, including the relentless threat of viral infections that can severely impact crop health and yield [1]. Cotton viruses, transmitted primarily by insect vectors such as aphids and whiteflies, pose a significant risk to cotton cultivation. Some of the most common cotton viruses include Cotton leaf curl virus (CLCuV), Cotton leaf crumple virus (CLCrV), and Cotton boll rot virus (CBRV). These viruses can cause symptoms such as leaf curling, stunted growth, yellowing, and reduced boll formation, ultimately leading to reduced yields and economic losses [2]. Given its economical importance, cotton has been the focus of intensive virology studies worldwide [1]. Even though Argentina is one important cotton producer, the study of pathogens in general, and viruses in particular associated to this crop are limited [3].

Geminiviruses (family *Geminiviridae*) are insect transmitted, small non-enveloped viruses, with circular single-stranded DNA genomes (2.5-5.2 kbps), which are encapsidated in quasi-icosahedral geminated virions of 22-38 nm size [4]. Geminiviruses are significant crop pathogens and cause important damages to economically relevant vegetables in many tropical and subtropical areas of the world [4]. Begomoviruses have mono or bipartite genomes and are transmitted by whiteflies (*Bemisia tabaci*). Genus *Begomovirus* currently harboring 409 species is the largest genus of plant infecting viruses, some of them causing severe diseases in vegetables and ornamental crops [4].

A cotton sample consisting of a *G. hirsutum* plant showing mosaic symptoms collected on May of 2015 from Argentina was subjected to high-throughput sequencing (HTS). To this end, total RNA from a symptomatic leaf cotton sample was purified using the RNAqueous small scale phenol-free total RNA isolation kit (Ambion), depleted of ribosomal RNA with RiboZero (Illumina, USA) and subjected to HTS in an Illumina Novaseq 6000 instrument generating 70,593,063 paired reads (150 nt long). These reads were trimmed and filtered with the Trimmomatic tool, as implemented in http://www.usadellab.org/cms/?page=trimmomatic, and the resulting 68,325,552 reads were mapped with Bowtie2 available at http://bowtie-bio.sourceforge.net/bowtie2/ using standard parameters against the *G. hirsutum* genome (assembly v2.1) available at https://www.ncbi.nlm.nih.gov/datasets/genome/GCF_007990345.1/. The resulting 34,174,101 unaligned paired reads were assembled *de novo* with Trinity <u>v2.9.0 release</u> with standard parameters. The obtained transcripts were subjected to bulk local Blastx searches (E-value <1e^−5^) against a complete NCBI Refseq of virus proteins database retrieved at ftp://ftp.ncbi.nlm.nih.gov/refseq/release/viral/viral.1.protein.faa.gz. Only two contigs of 1,896 and 2,386 nt long obtained significant hits to virus proteins, more specifically to the replication associated protein (REP) of Sida mosaic Bolivia virus 2 (SiMBoV2, E-value = 0.0, Identity 86.03%, GenBank accession number YP_004207815.1), and to the nuclear shuttle protein (NSP) of SiMBoV2, E-value = 5e-153, Identity 83.98%, GenBank accession number YP_004207829.1), respectively. To further extend these transcripts the filtered reads were iteratively mapped to these contigs, dubbed putDNA-A and putDNA-B, using the Geneious v8.1.9 platform (Biommaters, USA) map-to-reference tool with low sensitivity parameters as described elsewhere [5, 6]. Eventually, both sequences were widely extended and supported by more than two thousand reads each (putDNA-A: 2,680 nt long, supported by 287,483 reads, mean coverage = 14,618X and putDNA-B: 2,645 nt long, supported by 307,080 reads, mean coverage = 16,363X). Interestingly, read mapping indicated a circular nature of both sequences, which was confirmed by identifying overlapping reads at all positions, and thus a continuum of virus reads supported circularity at all sequence positions.

Blastn analysis of sequences putDNA-A and putDNA-B using default parameters showed the greatest nucleotide sequence identity to DNA component A of an isolate of SiMBoV2 of *Sida micrantha* from Bolivia (HM585443.1, coverage 100%, E-value 0.0, Identity 84.89%), and to DNA component B of an isolate of SiMBoV2 of chia (*Salvia hispanica*) from Argentina (KJ742422.1, coverage 100%, E-value 0.0, Identity 78.61%), respectively (**Table 1**; **Supplementary Table 1**). A rapid search of the sequences for open reading frames (ORFs) greater than 150 nt in size with the program ORF Finder (https://www.ncbi.nlm.nih.gov/orffinder/) indicated that the sequence architecture of putDNA-A and putDNA-B resembled that of SiMBoV2 New World bipartite begomovirus genome. That is, putDNA-A presented one ORF in the virion sense (V) and four in the complementary sense (C), and putDNA-B presented one ORF in the virion sense (V) and one in the complementary sense (C) (**Figure 1**). ORF AV1 of putDNA-A is predicted to encode a 249 aa coat protein (CP) sharing a 94.8% aa pairwise identity (PI) to that of SiMBoV2, and presenting a Gemini_coat domain (pfam00844, coordinates 13-249 aa, E-value = 1.21e-98) according to the NCBI-CDD conserved domain database available at https://www.ncbi.nlm.nih.gov/Structure/cdd/wrpsb.cgi. To confirm these putative virus sequences to cotton we designed specific primers targeting the CP CDS of putDNA-A. Total RNA was independently purified from the leaf sample of symptomatic cotton employed for the HTS library and of an apparently healthy plant sampled also in May 2015, using the same conditions as described above. The synthesis of cDNA from 3 μg of total RNA was carried out using SuperScript III reverse transcriptase and random primers (Invitrogen). The resulting cDNAs were amplified by standard PCR with primers CoMV-Fw: GTTAAGTGGGCTCGACACT and CoMV-Rv: GTTCGAGCTTCTGGGAGGAG targeting the AV1 and part of the AC3 coding sequence of DNA-A, which we designed using PrimerBLAST (https://www.ncbi.nlm.nih.gov/tools/primer-blast/), to avoid *G. hirsutum* endogenous targets. PCR reactions included Platinum Taq DNA Polymerase (Invitrogen) and used a program consisting of 35 cycles of 30 s at 94 °C, 30 s at 55 °C, 1 min 30 s at 72 °C, and a final elongation step of 2 min at 72 °C. The PCR reactions were run on standard agarose gel electrophoresis. Only the sample from the symptomatic leaf showed a PCR product of the expected size (973 nt) on the gels (**Supplementary Figure 4**). In addition, the PCR product was purified by ExoSAP-IT™ PCR Product Cleanup Reagent and subjected to Sanger sequencing, which confirmed with a 100% identity to putDNA-A target region of the amplicons amplified from the independent leaf DNA symptomatic sample. Thus, we hypothesized that the circular assembled sequences correspond to genome component A and B a new virus linked to *G. hirsutum* that we tentatively dubbed cotton mosaic virus (CoMV). To entertain this hypothesis, we advanced in the structural and functional annotation of the proposed virus sequences. When comparing DNA-A and B of CoMV by MUSCLE sequence alignments we detected a common region (CR), 201 nt long sharing a 95.5% PI. This bipartite begomovirus typical region suggests that the DNA components correspond to the same virus. Further inspection of this CR showed three repeated iteron like regions “TGGTACTCA” (two in the virion and one in the complementary strand) upstream of a TATA box and the canonical nonanucleotide sequence “TAATATT/AC” that marks the origin of virion-strand DNA replication, which was designated as position 1 at the nick region. DNA foldings of the CR using the MFOLD server implemented at http://unafold.rna.albany.edu/?q=mfold/DNA-Folding-Form suggested that, as expected, the nonanucleotide sequence is accessible to the REP protein in the loop of a well-supported stem-loop (**Supplementary Figure 1**). Besides the aforementioned AV1 CDS, DNA-A presented four ORFs in the complementary sense (AC1-AC4) (**Figure 1**). AC1 (coordinates 2,550-1,474 nt) is predicted to encode a 358 aa replication-associated protein (REP) sharing a 86% aa PI to that of SiMBoV2. The N-terminal region presented an Origin of replication-binding domain, RBD-like (SSF55464, coordinates 4-121 aa, E-value = 1.13e-57) as indicated by the SUPERFAMILY HMM search platform implemented at http://supfam.org/SUPERFAMILY/cgi-bin/hmm.cgi. In addition, the REP protein presented a Gemini_AL1 domain (pfam00799, coordinates 7-119 aa, E-value = 3.71e-64), a Gemini_AL1_M domain (pfam08283, coordinates 126-231 aa, E-value = 5.43e-46), followed by a REP associated domain (PD000736, coordinates 248-334 aa, E-value = 3e-17) according to the ProDom tool available at http://prodom.prabi.fr/. The overlapping ORF AC2 (coordinates 1,553-1,164 nt) encodes a 129 aa transcriptional regulator protein (TrAP/AL2) sharing a 86.8% aa PI to that of SiMBoV2, and presented a Gemini_AL2 domain (pfam01440, coordinates 1-124 aa, E-value = 1.67e-42). ORF AC3 (coordinates 1,417-1,019 nt) encodes a 132 aa replication enhancer protein (REn) sharing a 85.6% aa PI to that of SiMBoV2, and presented a Gemini_AL3 domain (pfam01407, coordinates 2-121 aa, E-value = 5.19e-50). Lastly, ORF AC4 (coordinates 2,399-2,136 nt), completely embedded within ORF AC1, encodes a 132 aa AC4 protein (REn) sharing a 77% aa PI to that of SiMBoV2, with a Gemini_C4 domain (pfam01492, coordinates 3-86 aa, E-value = 3.46e-35). On the other hand, the 2,645 nt long DNA-B, with two ORFS, encoded a nuclear shuttle protein (NSP) in virion ORF BV1 (505-1,275 nt coordinates) and a movement protein in ORF BC1 (coordinates 2,260-1,379 nt). The 256 aa NSP shared a 84% aa PI to that of SiMBoV2, and had a Gemini_coat domain (pfam00844, coordinates 7-256 aa, E-value = 1.70e-90). The 293 aa movement protein shared a 93.2% aa PI to that of SiMBoV2, and presented a Gemini_BL1 domain (pfam00845, coordinates 13-292 aa, E-value = 0) (**Figure 1**).

**Figure 1.**
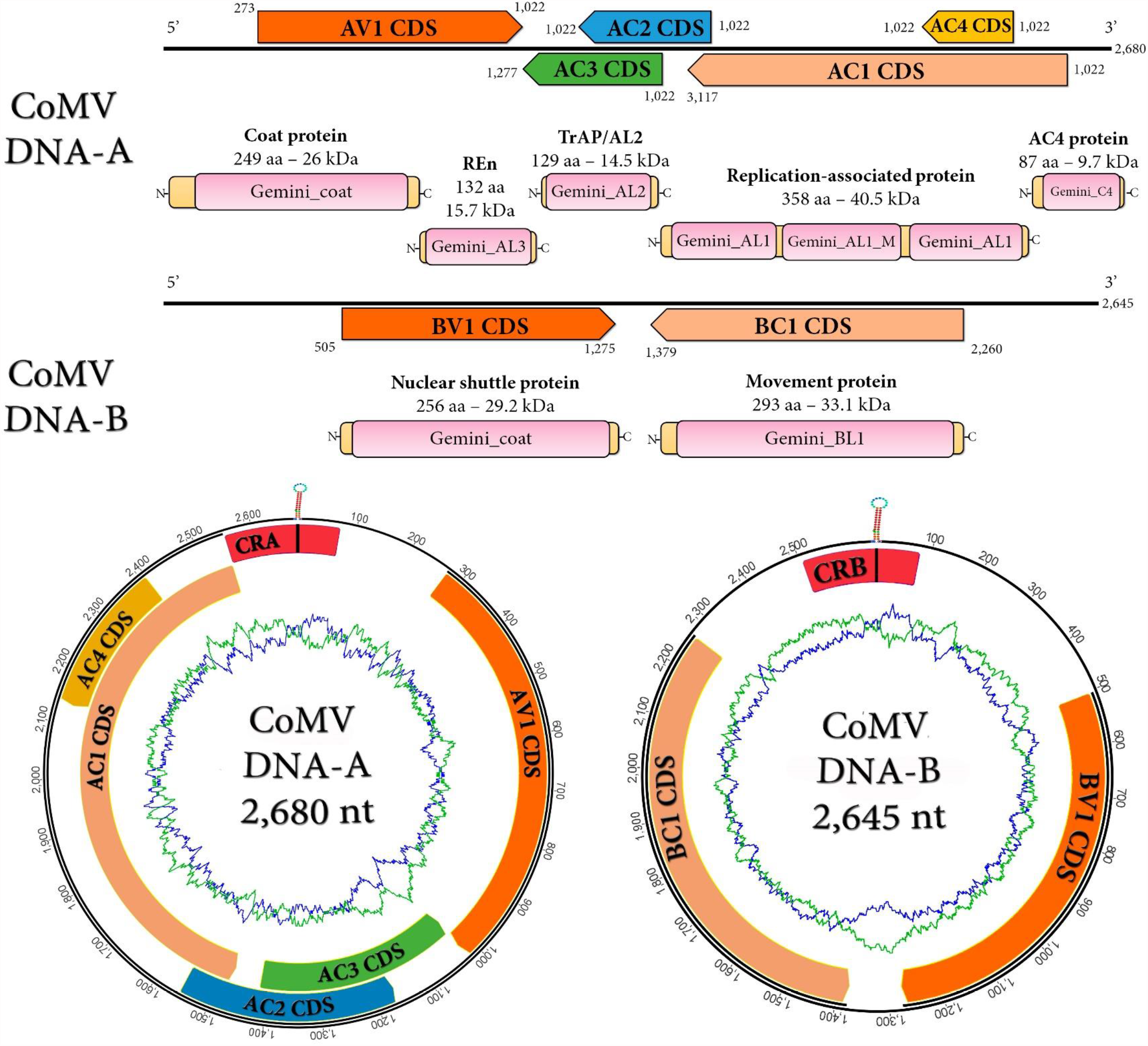
Molecular characterization of cotton mosaic virus (CoMV). Genome graph depicting predicted circular DNA components A and B of CoMV. Start and end coordinates of each gene product are indicated. Arrowed rectangles represent each ORF. Curved yellow rectangles represent each predicted proteins and conserved domains are shown in pink. In the lower panel the circular track indicates GC contant variation (GC blue, AT green). Abbreviations: AV1/BV1, virion strand ORF 1 of DNA-A or DNA-B; AC1-4/BC1, complementary strand ORF1-4 of DNA-A or DNA-B; Ren, replication enhancer protein; TrAP, transcriptional regulator protein; CRA/CRB, common region of DNA-A or DNA-B.

Given the clear evidence based on genomic architecture and structural annotation that the identified virus sequences corresponded to a New World bipartite begomovirus, we advanced with further genetic distance and evolutionary insights. These analyses were oriented to assess whether we could determine if the virus corresponded to a new strain of a previously reported virus such as SiMBoV2, or to a tentative member of a novel species of begomovirus. Firstly, we retrieved the 50 closest hits based on Blastn searches with standard parameters using as query complete DNA components A and B, and additionally African cassava mosaic virus (ACMV, NC_001467-NC_001468). As observed in **Table 1** and **Supplementary Table 1**, no virus sequences presented sequence identity higher than 84.89% (to DNA-A), or 78.61% (to DNA-B). Nevertheless, the current criteria for begomovirues recommended by the *Geminiviridae* Study Group of the International Committee on Taxonomy of Viruses indicates a threshold of <91% nt sequence identity for the complete DNA-A as a requisite for species demarcation [7], in the context of identities calculated from true pairwise sequence identities excluding gaps. Thus, as recommended, we employed the Sequence Demarcation Tool (SDT) v. 1.0 [8] with MUSCLE [9] alignment option to generate pairwise sequence comparisons. Our results confirm that the identified virus sequences are well below this <91% nt sequence identity threshold, being SiMBoV2 (HM585443) the highest identity virus with 85.9% PI for DNA-A, and 80.1% PI with SiMBoV2 (KJ742422) DNA-B (**Supplementary Figure 2, Supplementary Table 2**). In this scenario, we propose that the detected sequences correspond to a new virus, a tentative member of a novel species, which we suggest the name “cotton mosaic virus” (CoMV). To complement these findings we generated phylogenetic insights based on multiple genome alignments of both DNA-A and B components using MAFFT v7.017 https://mafft.cbrc.jp/alignment/server/ (1PAM/k2 scoring matrix), using E-INS-i as the best-fit model. The aligned genomes were used as input for maximum likelihood phylogenetic trees using FastTree 2.1.5 as implemented in http://www.microbesonline.org/fasttree/ (best-fit model = JTT-Jones-Taylor-Thorton with single rate of evolution for each site = CAT) computing local support values with the Shimodaira-Hasegawa test (SH) and 1,000 tree resamples, the trees were rooted at ACMV. Our results suggest that CoMV unequivocally clusters within genus *Begomovirus*, in a separated clade with affinity to SiMBoV2 and other Sida sp., Solanaceae, Salvia and Okra viruses (**Figure 2, Supplementary Figure 3**). It is worth mentioning that CoMV highest identity viruses correspond to isolates of viruses from Sida sp. which as cotton, is a member of the very same sub-family of plants *Malvoideae* (*Malvaceae*). In addition, the high identify viruses were detected in our region, that is mostly Brazil and Argentina (**Table 1, Supplementary Table 1)** and that there is an apparent clustering of viruses according to geographic distribution on phylogenetic trees **(Figure 2)**. This is in line with a suggested evolutionary path of divergence through isolation [10] as a consequence of the incapacity of whiteflies, the most plausible vectors, to fly long distances.

**Figure 2.**
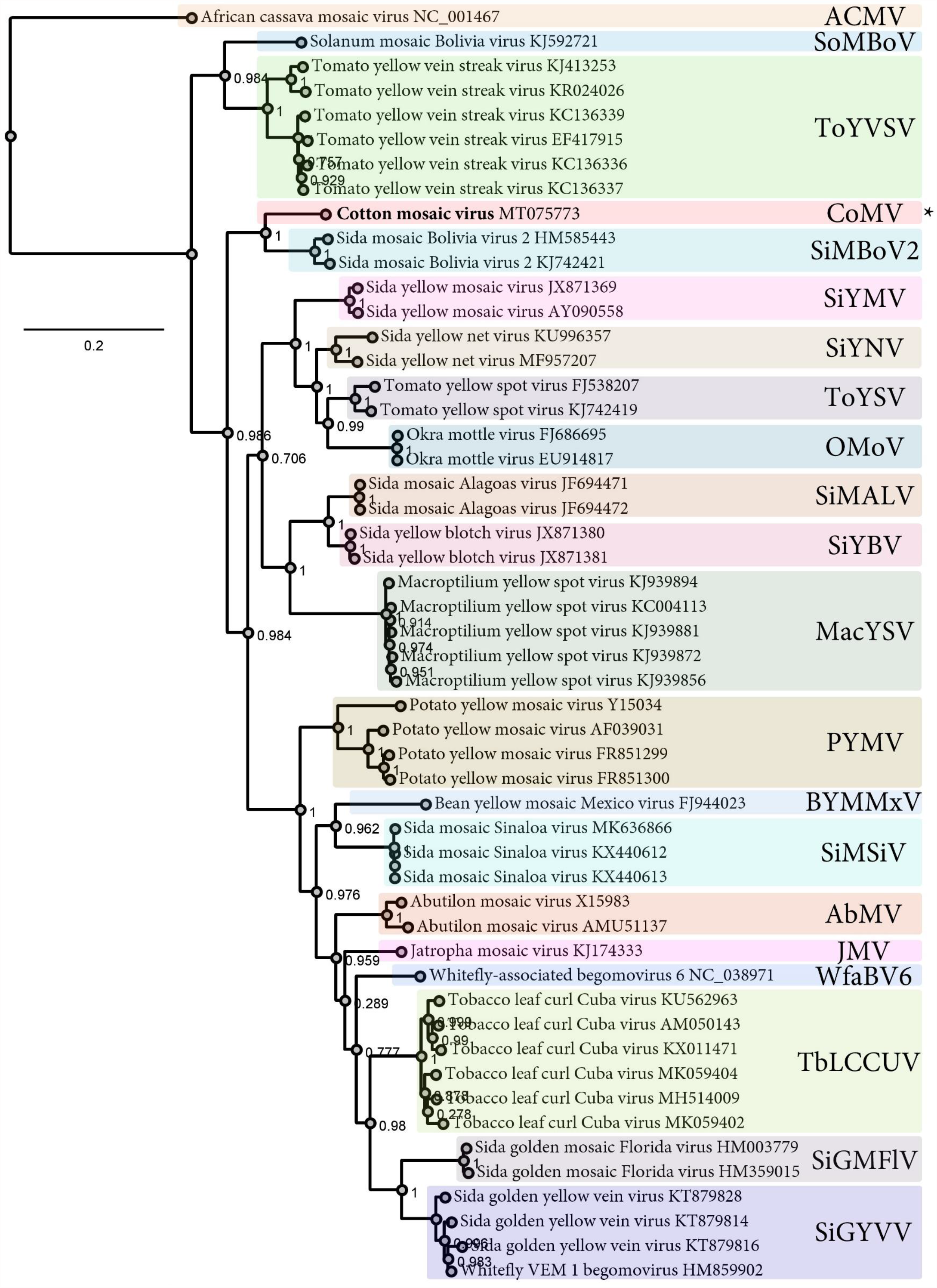
Phylogenetic insights of cotton mosaic virus (CoMV) based on MAFFT alignments and maximum likelihood trees generated with DNA-A of CoMV and reported begomoviruses. The scale bar indicates the number of substitutions per site. Node labels indicate FastTree support values. The tree is rooted at ACMV. Genbank Accession numbers are indicated following the virus names. Viruses corresponding to the same species are demarcated with colored rectangles. Standard ICTV virus abbreviations are shown on the right.

While this symptomatic cotton plant was examined our very same group [11] was assessing the viral landscape of grapevine samples. We started surveying *Vitis vinifera* samples from several regions and cultivars, representing the most significant wine cultivars from the main viticulture areas of Argentina. A pooled sample consisting of *V. vinifera* cv Torrontes from Mendoza province and four criollas varieties were subjected to high-throughput sequencing (HTS). The resulting transcripts assembled from the RNAseq data were assessed against viruses, and we found evidence of several well-known grapevine viruses. Strikingly, the very same begomovirus described here was also found in the *V. vinifera* RNA data. Our unawareness of many technical issues and our rush to communicate these findings, resulted in the unfortunate preprinting of that data linking the begomovirus to grapevine after an indirect confirmation of the findings based on PCR amplification and a restriction assay pattern consistent with the expected *in silico* data [11]. After numerous failed attempts to replicate our PCR amplifications and generate sanger data consistent with this begomovirus in grapevine we dabbled in this cotton project, assessed the obtained RNAseq data and found the exact same begomovirus. After we confirmed the detection of this virus in cotton and various efforts to understand what happened in grapevine we eventually found out that the samples which were separately and in different dates purified and processed to generate sequencing libraries were fortuitously sequenced at a provider in the same experiment and with spill over from the cotton sample to the grapevine sample probably by index hopping [12, 13]. That is the grapevine RNAseq data had a significantly small proportion of cotton data, which was fairly enough that we detected the begomovirus, which was a new one at that time, and we were able to assemble the complete genome with a relatively robust depth (97-109x mean coverage). Even though we routinely work with RNAseq data from numerous sources including third party publicly available data and we usually detect inconsistencies readily pointing to deficient data (human viruses in plant RNA samples, fungal viruses in insect RNA samples, etc.), this scenario raises a concern of the difficulties when dealing with new virus detections on metatranscriptomic data which may harbor more than one “plausible” host for the detected virus and the technical issues to demarcate the *bona fide* provenance of the HTS resource. That being said, we apologize to all researchers from the grapevine community who may have lost time and resources trying to replicate our findings, specially for taking so much time to revise this work.

All in all, our results suggest that a novel New World bipartite begomovirus is associated to cotton. As of today, we have no additional information on the infectivity, host range, geographical distribution or biology of CoMV. Nevertheless, we believe that this report, the associated sequences, and the newly developed detection specific-primers would aid in further characterizing the genetic variability of this virus which would eventually redound in an assessment of the prevalence of this agent in the region and worldwide. To our knowledge, this is the first report of a this new begomovirus linked to cotton, which highlights the diversity of viruses able to infect this economically important crop.

## Data availability

The genome sequences reported here are available at GenBank: CoMV-DNA-A (MT075773), CoMV-DNA-B (MT075774) and as Supplementary Material of this article.

## Acknowledgements

We thank P. Schubert for helpful discussions and insightful comments.

## Ethical approval

This article does not contain any studies with human participants or animals performed by any of the authors.

## Figure Legends

**Supplementary Figure 1.** (**A**) MUSCLE sequence alignments of DNA-A and B of CoMV common regions (CR), sharing a 95.5% PI. The CR presented three repeated iteron like regions “TGGTACTCA” (shown in green) upstream of a TATA box (in red) and the canonical nonanucleotide sequence “TAATATT/AC” (in purple) flanked by complementary regions (in red). (**B**) DNA folding of a sub region of the CR using the MFOLD server showing a well-supported stem-loop flanking the typical nonanucleotide sequence in the loop. The bases are colored by base pair probability from blue to red. Nonanuclotide sequence is demarcated with a colored rectangle. Nick region is indicated with a dotted line.

**Supplementary Figure 2.** Pairwise sequence comparisons plots of CoMV DNA-A (**A**) or DNA-B (**B**) and reported begomovirus including the 50 closest Blastn hits using as query the complete genome complement and ACMV. The plots were generated with the Sequence Demarcation Tool (SDT) v. 1.0 with MUSCLE alignment option. SDT matrix scores are available in **Supplementary Table 2**.

**Supplementary Figure 3.** Phylogenetic insights of cotton mosaic virus (CoMV) based on MAFFT alignments and maximum likelihood trees generated with DNA-A of CoMV and reported begomoviruses. The scale bar indicates the number of substitutions per site. Node labels indicate FastTree support values. The tree is rooted at ACMV. Genbank Accession numbers are indicated following the virus names. Viruses corresponding to the same species are demarcated with colored rectangles. Standard ICTV virus abbreviations are shown on the right.

**Supplementary Figure 4.** PCR confirmation of virus sequence. Specific primers were designed to detect CoMV DNA-A, with primers CoMV-Fw: GTTAAGTGGGCTCGACACT and CoMV-Rv: GTTCGAGCTTCTGGGAGGAG targeting the AV1 and part of the AC3 coding sequence of CoMV DNA-A. The only sample which generated an amplification product (ca. 973 nt) corresponded to the DNA extraction of the symptomatic cotton sample. The PCR product was purified and Sanger sequenced showing a 100% identity with the expected target region.

