## Supplementary material for "A New World begomovirus infecting Cotton in Argentina": Table 1

**Table 1**. Virus sequences with highest identity with Cotton mosaic virus corresponding to DNA component A based on Blastn searches.

| **Genbank Accession #** | **Virus**  **species** | **Isolate** | **Plant**  **Host** | **Origin** | **Total Score** | **Query Cover** | **E value** | **Per. Ident** |
| --- | --- | --- | --- | --- | --- | --- | --- | --- |
| [HM585443.1](https://www.ncbi.nlm.nih.gov/nucleotide/HM585443.1?report=genbank&log$=nucltop&blast_rank=1&RID=2B7B64RZ016) | *Sida mosaic Bolivia virus 2* | SiMBoV2 | *Sida micrantha* | Bolivia | 3021 | 100% | 0.0 | **84.89%** |
| [KJ742421.1](https://www.ncbi.nlm.nih.gov/nucleotide/KJ742421.1?report=genbank&log$=nucltop&blast_rank=2&RID=2B7B64RZ016) | *Sida mosaic Bolivia virus 2* | SM1 | *Salvia hispanica* | Argentina | 3012 | 100% | 0.0 | **84.84%** |
| [FJ538207.1](https://www.ncbi.nlm.nih.gov/nucleotide/FJ538207.1?report=genbank&log$=nucltop&blast_rank=3&RID=2B7B64RZ016) | *Tomato yellow spot virus* | ARGbean | *Phaseolus vulgaris* | Argentina | 2570 | 91% | 0.0 | **83.45%** |
| [MF957207.1](https://www.ncbi.nlm.nih.gov/nucleotide/MF957207.1?report=genbank&log$=nucltop&blast_rank=4&RID=2B7B64RZ016) | *Sida yellow net virus* | SaoFidelis_RJ | *Passiflora edulis* | Brazil | 2507 | 93% | 0.0 | **82.42%** |
| [JX871369.1](https://www.ncbi.nlm.nih.gov/nucleotide/JX871369.1?report=genbank&log$=nucltop&blast_rank=5&RID=2B7B64RZ016) | *Sida mosaic Bolivia virus 2* | BR:Vic1:10 | *Sida santaremensis* | Brazil | 2511 | 93% | 0.0 | **82.48%** |
| [KU996357.1](https://www.ncbi.nlm.nih.gov/nucleotide/KU996357.1?report=genbank&log$=nucltop&blast_rank=6&RID=2B7B64RZ016) | *Sida yellow net virus* | RJ028 | *Solanum lycopersicum* | Brazil | 2489 | 92% | 0.0 | **82.35%** |
| [MF957205.1](https://www.ncbi.nlm.nih.gov/nucleotide/MF957205.1?report=genbank&log$=nucltop&blast_rank=7&RID=2B7B64RZ016) | *Sida yellow net virus* | Paragominas_PA | *Passiflora edulis* | Brazil | 2486 | 93% | 0.0 | **82.18%** |
| [JX871376.1](https://www.ncbi.nlm.nih.gov/nucleotide/JX871376.1?report=genbank&log$=nucltop&blast_rank=8&RID=2B7B64RZ016) | *Sida yellow net virus* | BR:Vic2:10 | *Sida micrantha* | Brazil | 2476 | 89% | 0.0 | **83.05%** |
| [JX871380.1](https://www.ncbi.nlm.nih.gov/nucleotide/JX871380.1?report=genbank&log$=nucltop&blast_rank=9&RID=2B7B64RZ016) | *Sida yellow blotch virus* | BR:Rla1:10 | *Sida urens* | Brazil | 2438 | 92% | 0.0 | **81.91%** |
| [KJ742419.1](https://www.ncbi.nlm.nih.gov/nucleotide/KJ742419.1?report=genbank&log$=nucltop&blast_rank=10&RID=2B7B64RZ016) | *Tomato yellow spot virus* | TO1 | *Salvia hispanica* | Argentina | 2438 | 90% | 0.0 | **82.18%** |
| [JF694472.1](https://www.ncbi.nlm.nih.gov/nucleotide/JF694472.1?report=genbank&log$=nucltop&blast_rank=11&RID=2B7B64RZ016) | *Sida mosaic Alagoas virus* | BgV02A.1.C61 | *Sida sp.* | Brazil | 2429 | 89% | 0.0 | **82.41%** |
| [JF694471.1](https://www.ncbi.nlm.nih.gov/nucleotide/JF694471.1?report=genbank&log$=nucltop&blast_rank=12&RID=2B7B64RZ016) | *Sida mosaic Alagoas virus* | BgV02A.1.C59 | *Sida sp.* | Brazil | 2429 | 89% | 0.0 | **82.41%** |
| [EU914817.1](https://www.ncbi.nlm.nih.gov/nucleotide/EU914817.1?report=genbank&log$=nucltop&blast_rank=13&RID=2B7B64RZ016) | *Okra mottle virus* | 6319 | *Abelmoschus esculentus* | Brazil | 2429 | 88% | 0.0 | **82.73%** |
| [FJ686695.1](https://www.ncbi.nlm.nih.gov/nucleotide/FJ686695.1?report=genbank&log$=nucltop&blast_rank=14&RID=2B7B64RZ016) | *Okra mottle virus* | BR:Sag8:Soy:08 | *Glycine max* | Brazil | 2426 | 88% | 0.0 | **82.65%** |
| [AY090558.1](https://www.ncbi.nlm.nih.gov/nucleotide/AY090558.1?report=genbank&log$=nucltop&blast_rank=15&RID=2B7B64RZ016) | *Sida yellow mosaic virus* | [Brazil] | *Sida sp.* | Brazil | 2407 | 93% | 0.0 | **81.64%** |
| [JX871381.1](https://www.ncbi.nlm.nih.gov/nucleotide/JX871381.1?report=genbank&log$=nucltop&blast_rank=16&RID=2B7B64RZ016) | *Sida yellow blotch virus* | BR:Rla2:10 | *Sida urens* | Brazil | 2393 | 92% | 0.0 | **81.54%** |
| [EU914819.1](https://www.ncbi.nlm.nih.gov/nucleotide/EU914819.1?report=genbank&log$=nucltop&blast_rank=17&RID=2B7B64RZ016) | *Okra mottle virus* | 6328 | *Abelmoschus esculentus* | Brazil | 2410 | 88% | 0.0 | **82.58%** |
| [KX348179.1](https://www.ncbi.nlm.nih.gov/nucleotide/KX348179.1?report=genbank&log$=nucltop&blast_rank=18&RID=2B7B64RZ016) | *Tomato yellow spot virus* | BR:Pab1058:11 | *Leonurus sibiricus* | Brazil | 2402 | 88% | 0.0 | **82.24%** |
| [KJ174333.1](https://www.ncbi.nlm.nih.gov/nucleotide/KJ174333.1?report=genbank&log$=nucltop&blast_rank=19&RID=2B7B64RZ016) | *Jatropha mosaic virus* | DO:Aguacatico | *Jatropha sp.* | Do. Republic | 2267 | 84% | 0.0 | **82.22%** |
| [KC136337.1](https://www.ncbi.nlm.nih.gov/nucleotide/KC136337.1?report=genbank&log$=nucltop&blast_rank=20&RID=2B7B64RZ016) | *Tomato yellow vein streak virus* | CL:VS-B4:To:2012 | *Solanum lycopersicum* | Chile | 2206 | 94% | 0.0 | **79.35%** |
