## Supplementary figures and images for "A New World begomovirus infecting Cotton in Argentina"

### Supplementary Figure 1

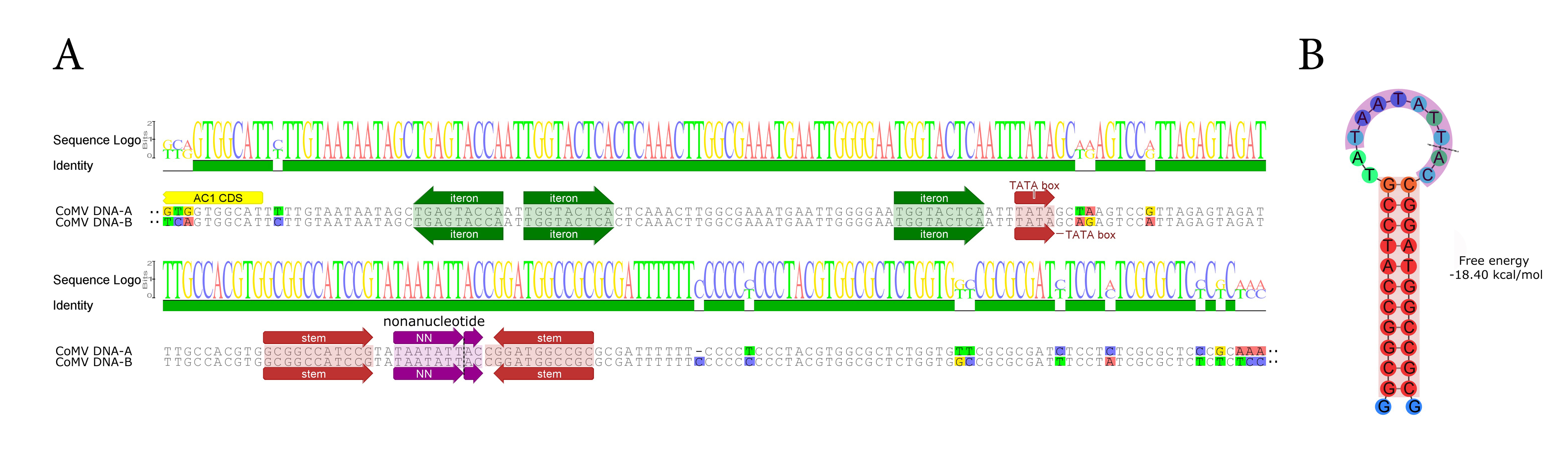

### Supplementary Figure 2

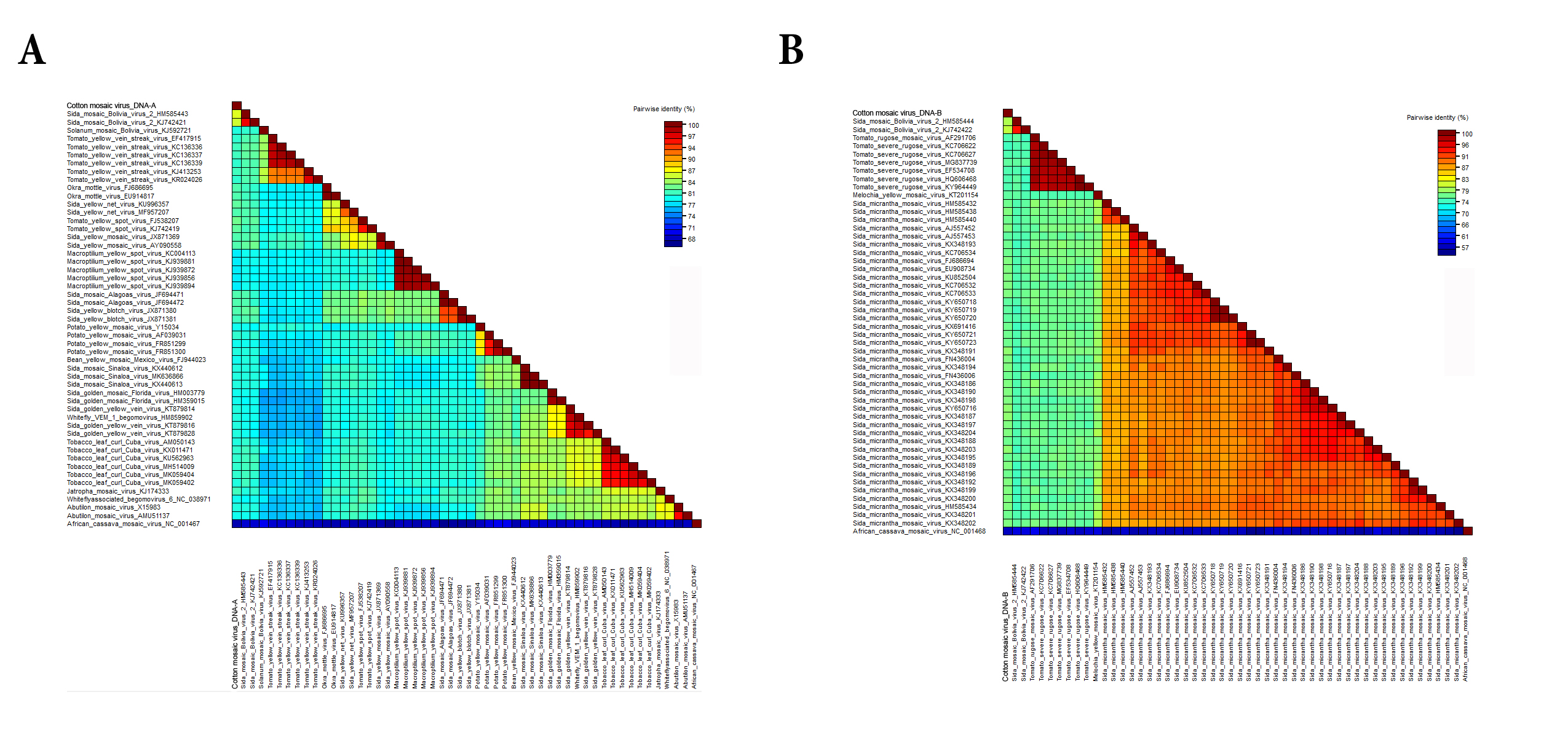

### Supplementary Figure 3

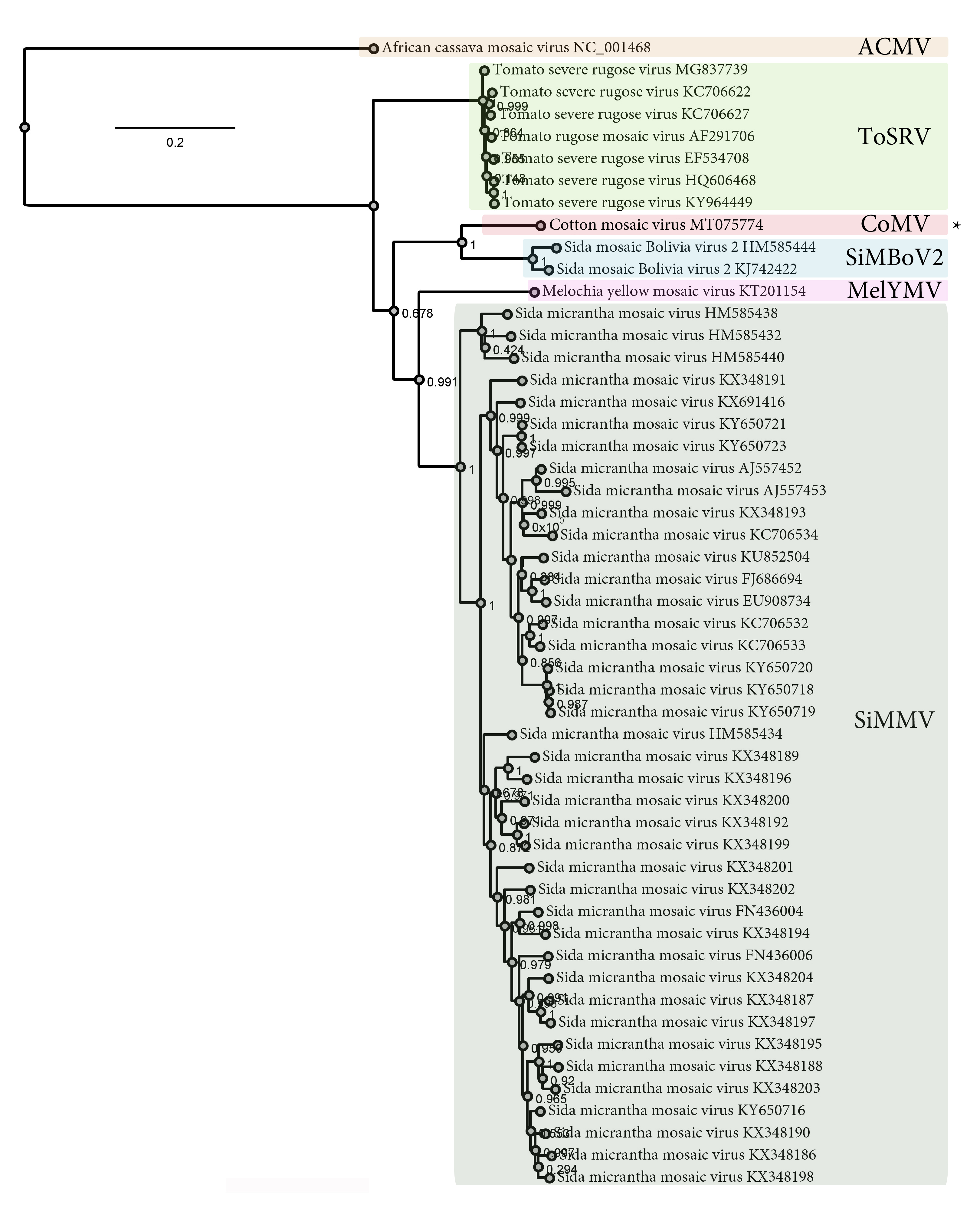

### Supplementary Figure 4

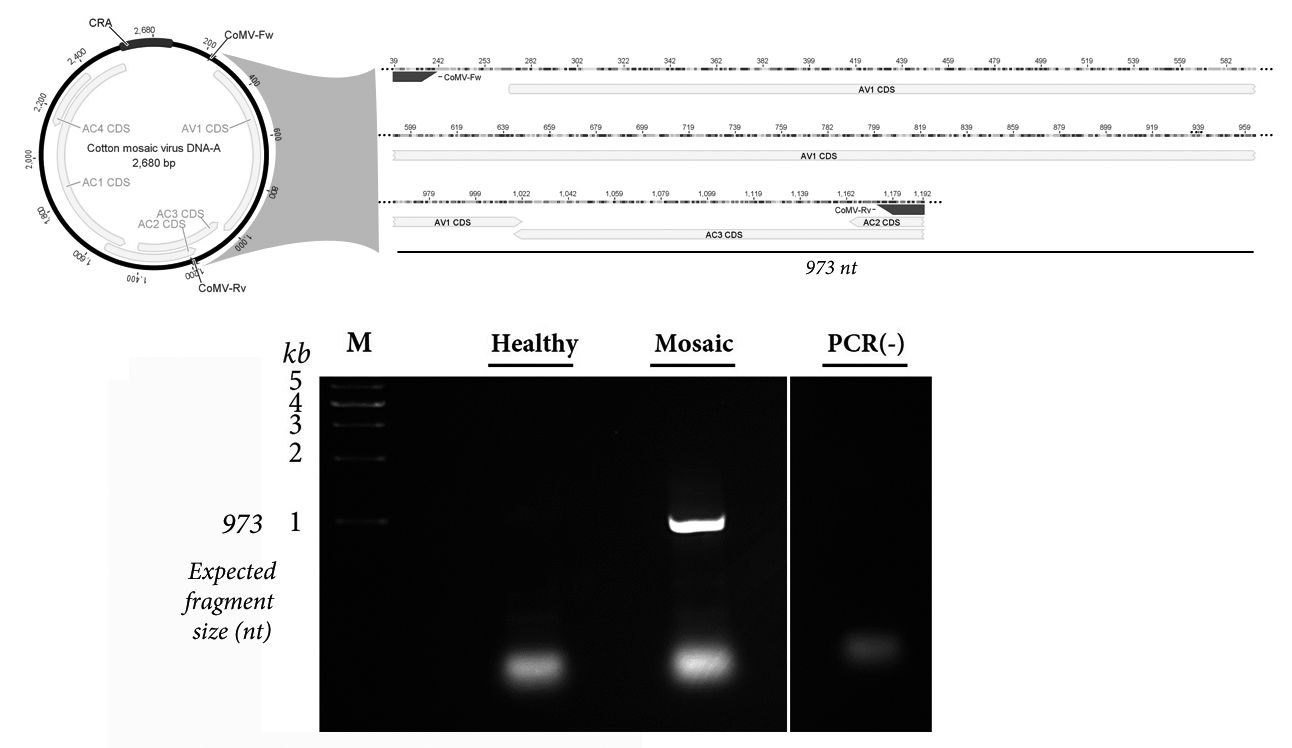
