## Supplementary Table 1 for "A New World begomovirus infecting Cotton in Argentina"

**Supplementary Table 1**. Virus sequences with highest identity with cotton mosaic virus corresponding to DNA component B based on Blastn searches.

| **Genbank Accession #** | **Virus**  **species** | **Isolate** | **Plant**  **Host** | **Origin** | **Total Score** | **Query Cover** | **E value** | **Per. Ident** |
| --- | --- | --- | --- | --- | --- | --- | --- | --- |
| [KJ742422.1](https://www.ncbi.nlm.nih.gov/nucleotide/KJ742422.1?report=genbank&log$=nucltop&blast_rank=1&RID=2BAY4Z91014) | *Sida mosaic Bolivia virus 2* | SM1 | *Salvia hispanica* | Argentina | 2183 | 100% | 0.0 | **78.61%** |
| [HM585444.1](https://www.ncbi.nlm.nih.gov/nucleotide/HM585444.1?report=genbank&log$=nucltop&blast_rank=2&RID=2BAY4Z91014) | *Sida mosaic Bolivia virus 2* | SiMBoV2 | *Sida micrantha* | Bolivia | 2117 | 100% | 0.0 | **78.10%** |
| [KX348204.1](https://www.ncbi.nlm.nih.gov/nucleotide/KX348204.1?report=genbank&log$=nucltop&blast_rank=3&RID=2BAY4Z91014) | *Sida micrantha mosaic virus* | BR:Gua1120:11 | *Sida sp.* | Brazil | 1861 | 84% | 0.0 | **79.57%** |
| [HM585434.1](https://www.ncbi.nlm.nih.gov/nucleotide/HM585434.1?report=genbank&log$=nucltop&blast_rank=4&RID=2BAY4Z91014) | *Sida micrantha mosaic virus* | SimMV-MGS2:07-Bo | *Sidastrum micranthum* | Bolivia | 1832 | 82% | 0.0 | **79.54%** |
| [KX348187.1](https://www.ncbi.nlm.nih.gov/nucleotide/KX348187.1?report=genbank&log$=nucltop&blast_rank=5&RID=2BAY4Z91014) | *Sida micrantha mosaic virus* | BR:Jan1128:11 | *Sida sp.* | Brazil | 1827 | 84% | 0.0 | **79.26%** |
| [KX348186.1](https://www.ncbi.nlm.nih.gov/nucleotide/KX348186.1?report=genbank&log$=nucltop&blast_rank=6&RID=2BAY4Z91014) | *Sida micrantha mosaic virus* | BR:Rea662:10 | *Sida sp.* | Brazil | 1806 | 84% | 0.0 | **79.05%** |
| [FN436004.1](https://www.ncbi.nlm.nih.gov/nucleotide/FN436004.1?report=genbank&log$=nucltop&blast_rank=7&RID=2BAY4Z91014) | *Sida micrantha mosaic virus* | BR:MGdosul:1:2007 | *Sida rhombifolia* | Brazil | 1814 | 84% | 0.0 | **78.90%** |
| [KX348202.1](https://www.ncbi.nlm.nih.gov/nucleotide/KX348202.1?report=genbank&log$=nucltop&blast_rank=8&RID=2BAY4Z91014) | *Sida micrantha mosaic virus* | BR:Saa832:10 | *Sida sp.* | Brazil | 1820 | 84% | 0.0 | **79.20%** |
| [KY650716.1](https://www.ncbi.nlm.nih.gov/nucleotide/KY650716.1?report=genbank&log$=nucltop&blast_rank=9&RID=2BAY4Z91014) | *Sida micrantha mosaic virus* | BR:Lon:OSP2:5 | *Oxalis sp.* | Brazil | 1806 | 84% | 0.0 | **78.99%** |
| [KX348190.1](https://www.ncbi.nlm.nih.gov/nucleotide/KX348190.1?report=genbank&log$=nucltop&blast_rank=10&RID=2BAY4Z91014) | *Sida micrantha mosaic virus* | BR:Sad799:10 | *Sida sp.* | Brazil | 1791 | 84% | 0.0 | **78.89%** |
| [KX348192.1](https://www.ncbi.nlm.nih.gov/nucleotide/KX348192.1?report=genbank&log$=nucltop&blast_rank=11&RID=2BAY4Z91014) | *Sida micrantha mosaic virus* | BR:Pan115:09 | *Sida sp.* | Brazil | 1800 | 82% | 0.0 | **79.30%** |
| [KX348197.1](https://www.ncbi.nlm.nih.gov/nucleotide/KX348197.1?report=genbank&log$=nucltop&blast_rank=12&RID=2BAY4Z91014) | *Sida micrantha mosaic virus* | BR:Mcr679:10 | *Sida sp.* | Brazil | 1790 | 84% | 0.0 | **78.96%** |
| [KX348198.1](https://www.ncbi.nlm.nih.gov/nucleotide/KX348198.1?report=genbank&log$=nucltop&blast_rank=13&RID=2BAY4Z91014) | *Sida micrantha mosaic virus* | BR:Mar755:10 | *Sida sp.* | Brazil | 1792 | 85% | 0.0 | **78.43%** |
| [HM585438.1](https://www.ncbi.nlm.nih.gov/nucleotide/HM585438.1?report=genbank&log$=nucltop&blast_rank=14&RID=2BAY4Z91014) | *Sida micrantha mosaic virus* | SimMV-rho[Bo:CF1:07] | *Sida rhombifolia* | Bolivia | 1787 | 88% | 0.0 | **78.27%** |
| [KX348199.1](https://www.ncbi.nlm.nih.gov/nucleotide/KX348199.1?report=genbank&log$=nucltop&blast_rank=15&RID=2BAY4Z91014) | *Sida micrantha mosaic virus* | BR:Cha822:10 | *Sida sp.* | Brazil | 1786 | 82% | 0.0 | **79.23%** |
| [KX348189.1](https://www.ncbi.nlm.nih.gov/nucleotide/KX348189.1?report=genbank&log$=nucltop&blast_rank=16&RID=2BAY4Z91014) | *Sida micrantha mosaic virus* | BR:Tap895:10 | *Sida sp.* | Brazil | 1775 | 84% | 0.0 | **78.87%** |
| [KX348200.1](https://www.ncbi.nlm.nih.gov/nucleotide/KX348200.1?report=genbank&log$=nucltop&blast_rank=17&RID=2BAY4Z91014) | *Sida micrantha mosaic virus* | BR:Smm876:10 | *Sida sp.* | Brazil | 1768 | 82% | 0.0 | **79.13%** |
| [KX348195.1](https://www.ncbi.nlm.nih.gov/nucleotide/KX348195.1?report=genbank&log$=nucltop&blast_rank=18&RID=2BAY4Z91014) | *Sida micrantha mosaic virus* | BR:Sam69:09 | *Sida sp.* | Brazil | 1761 | 84% | 0.0 | **78.55%** |
| [FN436006.1](https://www.ncbi.nlm.nih.gov/nucleotide/FN436006.1?report=genbank&log$=nucltop&blast_rank=19&RID=2BAY4Z91014) | *Sida micrantha mosaic virus* | BR:MGdosul:2:2007 | *Sida micrantha* | Brazil | 1757 | 84% | 0.0 | **78.61%** |
| [KX348194.1](https://www.ncbi.nlm.nih.gov/nucleotide/KX348194.1?report=genbank&log$=nucltop&blast_rank=20&RID=2BAY4Z91014) | *Sida micrantha mosaic virus* | BR:Cra556:10 | *Sida sp.* | Brazil | 1764 | 86% | 0.0 | **78.07%** |
